# Transcriptional Isoforms of NAD^+^ Kinase regulate oxidative stress resistance and melanoma metastasis

**DOI:** 10.1101/2023.02.02.526506

**Authors:** Graciela Cascio, Kelsey N. Aguirre, Kellsey P. Church, Riley O. Hughes, Leona A. Nease, Ines Delclaux, Marwa Zerhouni, Elena Piskounova

## Abstract

Metastasizing cancer cells encounter a multitude of stresses throughout the metastatic cascade. Oxidative stress is known to be a major barrier for metastatic colonization, such that metastasizing cancer cells must rewire their metabolic pathways to increase their antioxidant capacity. NADPH is essential for regeneration of cellular antioxidants and several NADPH-regenerating pathways have been shown to play a role in metastasis. We have found that metastatic melanoma cells have increased levels of both NADPH and NADP^+^ suggesting increased *de novo* biosynthesis of NADP^+^. *De novo* biosynthesis of NADP^+^ occurs through a single enzymatic reaction catalyzed by NAD^+^ kinase (NADK). Here we show that different NADK isoforms are differentially expressed in metastatic melanoma cells, with Isoform 3 being specifically upregulated in metastasis. We find that Isoform 3 is more potent in expanding the NADP(H) pools, increasing oxidative stress resistance and promoting metastatic colonization compared to Isoform 1. We have found that Isoform 3 is transcriptionally upregulated by oxidative stress through the action of NRF2. Together, our work presents a previously uncharacterized role of NADK isoforms in oxidative stress resistance and metastasis and suggests that NADK Isoform 3 is a potential therapeutic target in metastatic disease.

## Introduction

Oxidative stress limits metastasis^1-4^. Cancer cells, both in circulation and during metastatic colonization, experience high levels of oxidative stress and exhibit decreased levels of the cellular antioxidant, reduced glutathione (GSH)^5^. While most cancer cells do not survive the metastatic cascade, successful metastasizers are able to rewire their metabolism to upregulate antioxidant pathways and overcome oxidative stress by regenerating GSH from its oxidized form, GSSG. To reduce GSSG to GSH cells must utilize NADPH, in the process converting it to NADP^+^ (Supp Fig 1A). It has been shown previously that metastasizing cells rely on different NADPH regeneration pathways including one carbon metabolism and pentose phosphate pathway, to increase their antioxidant capacity. However, the process by which metastasizing cells reinforce their NADP(H) pool is unknown.

NADP^+^ is generated *de novo* by NAD^+^ kinase (NADK) by phosphorylating NAD^+6,7^. Due to a high cellular demand for reduction reactions, cells maintain a high NADPH/NADP^+^ ratio to ensure that these reactions are favorable^8^. Since NADP^+^ and NADPH are membrane impermeable cells have developed organelle-specific NADPH synthesis. In mammals, NADK is the cytoplasmic form of the enzyme, while NADK2 localizes to the mitochondria^9,10^. NADK contains a bifunctional NADP^+^ phosphatase/NAD kinase domain, while NADK2 contains a diacylglycerol kinase catalytic domain^11^. Phylogenetic analysis indicates that NADK and NADK2 have developed independently during evolution^11^. Presence of other NADKs in organelles such as peroxisomes and endoplasmic reticulum remains unknown. Recent studies demonstrated the importance of NADK in the antioxidant defense, while NADK2 has been mainly implicated in proline biosynthesis^12,13^.

Cytoplasmic NADK exists in three isoforms. Isoform 1 and 2 are the two longer isoforms that contain an autoinhibitory N-terminal domain^14^. It has been previously shown that both AKT and PKC regulate the activity of Isoform 1 through phosphorylation of three terminal serine residues: Ser44, Ser46 and Ser48^14,15^. This phosphorylation relieves the autoinhibition of NADK and stimulates NADP^+^ production. In contrast, Isoform 3 is a shorter isoform, generated through an alternative transcriptional start site with a unique N-terminal domain that does not have autoinhibitory activity or AKT/PKC phosphorylation sites. Consequently Isoform 3 has much higher enzymatic activity than Isoform 1 at steady state^14^. However, the differential role of NADK isoforms in NADP^+^ generation is unknown.

Here we examined the role of NADK isoforms in oxidative stress resistance and melanoma metastasis to address how NADK activity contributes to *de novo* NADP(H) generation as an antioxidant defense mechanism in metastasizing melanoma cells. We specifically focused on Isoforms 1 and 3 as Isoform 2 has the same regulatory domain as Isoform 1. We generated loss of function melanoma cell lines of these NADK Isoforms which allowed us to address how these differential activities impacted melanoma cell survival under oxidative stress and during metastasis. We demonstrated that metastasizing melanoma cells increase NADP(H) biosynthesis and increase levels of Isoform 3 compared to the primary tumor. Loss of Isoform 3 caused a greater increase in both cytoplasmic and mitochondrial oxidative stress levels than loss of Isoform 1, and sensitized cells to prooxidant treatment. Finally, loss of either Isoform 1 or 3 had no effect on the growth of subcutaneous tumors but lead to a significant decrease in metastasis. Interestingly, overexpression of Isoform 1 or 3 had no significant effect on primary tumor growth either, but overexpression of truncated Isoform 1 or Isoform 3 caused a much higher increase in metastatic burden in the organs of the mice compared to full-length Isoform 1. We found that transcriptional upregulation of Isoform 3 is driven by the transcription factor NRF2. Our findings suggest that metastasizing melanoma cells increase expression of NADK Isoform 3 to combat oxidative stress and increase their survival during the metastatic cascade. Our data indicate that increased levels of Isoform 3 promote melanoma cell extravasation and colonization of distant sites. Our findings identify NADK isoform 3 as a potential therapeutic target for metastatic disease.

## Methods

### Plasmid Construction

miR-E shRNA knockdown constructs were generated by the splashRNA program and ordered from Invitrogen as oligos. Hairpins were ligated (T4 Ligase, NEB) into SiREP lentiviral plasmid behind an SFFV promoter and iRFP fluorescent protein using XhoI and EcoRI sites. For overexpression constructs, the template plasmids for NADK isoforms were kindly provided by Gerta Hoxhaj and the coding sequences of NADK isoforms 1 and 3 for the rescue experiments were ordered as gene fragments (IDT) with an N-terminus Kozack sequence and 3xHA. Constructs were ligated (In-Fusion HD, Takara or T4 Ligase, NEB) into pLenti vector behind an EFS promoter using BamHI and NsiI sites. A cleavable P2A site links the protein expression to a fluorescent protein tag. CRISPR knockout sgRNA sequences were generated using Geneious cloning software and ordered as single stranded oligos from IDT. Vector backbones were kindly provided by Lukas Dow, and all plasmids were sequence verified.

### Lentiviral transduction and generation of stable cell lines

For virus production, 0.9μg of plasmid was combined with 1μg of packaging plasmids (0.4μg pMD2G and 0.6μg psPAX2) and transfected into HEK 293T cells using Polyjet (SignaGen) according to manufacturer’s instructions. Fresh media was added the following day and viral supernatants were collected 72hr after transfection and filtered through a 0.45μM filter. Approximately 1 million cells were infected with viral supernatant and 10μg/mL polybrene (Sigma-Aldrich). Cells then underwent antibiotic selection with puromycin or hygromycin (Inviogen) depending on the antibiotic resistance of each plasmid. For CRISPR knockout, cells stably-expressing Cas9 were infected with the gRNA construct. To generate single clones, cells were diluted and plated into a 10cm dish. Glass rings were used to isolate single cells and grow colonies. All single clones were verified by western blot analysis and Synthego genomic DNA sequencing.

### Real-Time PCR Quantification of mRNA

mRNA was extracted using a Direct-zol RNA Purification Kit (Zymo Research). Reverse transcription was performed using iScript cDNA synthesis kit (Rio-Rad) as described by manufacturer. cDNA was then diluted and used for qPCR analysis with SYBR Green PCR master mix (Thermo Fisher). IDT online primer design tool was used to generate qPCR primers and UCSC In-Silico PCR database was then used to verify human-specificity of primers. qPCR analysis was performed on an Applied Biosystems QuantStudio 6 Real-Time PCR System. All targets were normalized to Actin in a standard ∆∆Ct analysis.

### Analysis of cell survival under prooxidant treatment

Cells were plated at 10,000 cells/well in a white TC-treated 96 well plate (Corning). After 24 hours, cells were then treated with prooxidant H_2_O_2_ (Sigma-Aldrich). After 24 hours of treatment, CellTiter-Glo 2.0 (Promega) was used according to manufacturers instructions to quantify cell viability.

### Lentiviral transduction of human melanoma cells

A lentiviral construct with luciferase and dsRed (luc P2A dsRed) was used to label patient-derived melanoma cells as described in Piskounova et al. Nature (2015). Virus was produced as described above. For lentiviral transduction, 500,000 freshly dissociated melanoma cells were infected with viral supernatant and supplemented with 10 μg/mL polybrene (Sigma). The following day, the media was replaced and 48hrs post-infection cells were either injected subcutaneously into mice as bulk tumors or FACS sorted for positive infection.

### Flow cytometry of melanoma cells

All melanoma cells in this study stably express dsRed and luciferase so that melanoma cells could be distinguished by flow cytometry or bioluminescent imaging. When preparing cells for sorting by flow cytometry, cells were stained with antibodies against mouse CD45 (30-F11-VioletFluor, Tonbo), mouse CD31 (390-VioletFluor, eBiosciences), Ter119 (Ter-119-VioletFluor, Tonbo) and human HLA-A, -B, -C (BD Biosciences) to select live human melanoma cells and exclude mouse endothelial and hematopoietic cells. Antibody labelling was performed for 20 min on ice, followed by washing and centrifugation. Before sorting, cells were resuspended in staining medium (L15 medium containing bovine serum albumin (1 mg/mL), 1% penicillin/streptomycin, and 10mM HEPES, pH 7.4) containing 4’6-diamidino-2-phenylindole (DAPI; 5μg/mL; Sigma) to eliminate dead cells from sorting. Live human melanoma cells were isolated by flow cytometry by sorting cells that were positive for dsRed and HLA and negative for mouse CD45, CD31, Ter-119 and DAPI.

### Transplantation of melanoma cells

After sorting, cells were counted and resuspended in DMEM media with 50% high-protein Matrigel (product 354248, Corning). For a standard metastasis assay, 100 cells, for patientderived melanoma cells, or 50 for A375, were injected subcutaneously into the right flank of the mice. Tumor formation was evaluated by regular palpitation of the injection site and tumors were measured weekly until any tumor in the mouse cohort reached 2.5cm in its largest diameter. Mice were monitored weekly for signs of distress according to a standard body condition score or within 24hr of their tumors reaching 2.5 cm in diameter – whichever came first. These experiments were performed according to protocols approved by the Institutional Animal Care and Use Committee at Weill Cornell Medicine (protocol 2017-0033).

### Bioluminescence imaging

Mice injected subcutaneously with melanoma cells expressing luciferase were monitored until tumors reached 2.5cm in diameter. For bioluminescent imaging, mice were injected intraperitoneally with 100μL of DPBS containing D-luciferin monopotassium salt (40μg/mL, Goldbio) 5 min before imaging, followed by general anesthesia with isoflurane 2 min before imaging. IVIS Imaging System 200 Series (Caliper Life Sciences) with Living Image Software was used with the exposure time set to 10 seconds. After imaging the whole body, mice were euthanized and individual organs were dissected and imaged. The bioluminescent signal was quantified with ‘region of interest’ measurement tools in Living Image (Perkin Elmer) software.

After imaging, tumors and metastatic nodules were collected to perform protein and/or molecular analysis.

### Flow cytometric analysis of oxidative stress

Cells were plated in low attachment plates (3D system) or in TC-treated plates and treated with a prooxidant pyocyanin (Cayman). We stained the dissociated cells for 30 minutes at 37°C with 5 μM CellROX Green, or CellROX DeepRed in HBSS-free (Ca2+ and Mg2+ free) to assess mitochondrial and cytoplasmic oxidative stress.

### NADPH/NADP^+^ and NAD^+^/NADH measurement

For the mice results, subcutaneous tumors or metastatic nodules were surgically excised as quickly as possible after euthanizing the mice then melanoma cells were mechanically dissociated. For the melanoma cell lines, cell were plated the day before in 24 wells TC-treated plates. NADPH, NADP^+,^ NADH and NAD^+^ were measured using NADPH/NADP^+^ Glo-Assay (Promega) and NADH/NAD^+^ Glo-Assay (Promega) following the manufactures instructions.

Luminescence was measured using a Omega plate reader (BMG Labtech FLUOstar). Values were normalized to protein concentration, measured using The Pierce Rapid Gold BCA Protein Assay Kit (Thermo Fisher Scientific).

### Western Blot Analysis

Tissue or cell lines lysates were prepared using Triton lysis Buffer (Triton X-100 1%, 20mM Tris pH8, 137mM NaCl, 1mM EDTA, 10% glycerol, 1.5 mM MgCl_2_) supplemented with phenylmethylsulphonyl fluoride (Sigma-Aldrich), and 2 cocktails of protease and phosphatase inhibitor (Halt™, Fisher Scientific; Phosstop™-phosphatase inhibitor tablets, Roche). The Pierce Rapid Gold BCA Protein Assay Kit (Thermo Fisher Scientific) was used to quantify protein concentrations. Equal amounts of protein (15–30 μg) were separated on 4–20% Tris Glycine SDS gels (BioRad) and transferred to polyvinylidene difluoride membranes (BioRad). Membranes were blocked for 30 minutes at room temperature with 5% milk in TBS supplemented with 0.1% Tween20 (TBST) then incubated with primary antibodies overnight at 4°C. After incubating with horseradish peroxidase conjugated secondary antibodies (Cell Signaling Technology), membranes were developed using SuperSignal West Pico or Femto chemiluminescence reagents (Thermo Fisher Scientific). The following primary antibodies were used for western blot analyses: NADK (Cell Signaling Technologies; 55948S), NADK (Sigma-Aldrich; HPA048909), NADK2 (Abcam; ab181028), HA-Tag (Cell Signaling Technologies; 14031), β-Actin HRP Conjugate, (Cell Signaling Technologies; 12262).

### Cell culture

Melanoma cell line A375 was culture using DMEM (Corning; 10-017-CV) supplemented with 10% FBS and Penicillin: Streptomycin solution 100x (Corning; 45000-652).

### Statistical analysis

No statistical methods were used to predetermine sample size. The data in most figure panels represents several independent experiments performed on different days. Variation is always indicated using standard deviation. For analysis of statistical significance, we first tested whether there was homogeneity of variation across conditions (as required by ANOVA) using Levene’s test, or when only two conditions were compared, using the F-test. In cases where the variation was significantly different among conditions, we used a non-parametric Kruskal-Wallis test or a non-parametric Mann-Whitney test to assess significance of difference among populations and conditions. Usually variation did not significantly differ among conditions. Under those circumstances, two-tailed Student’s t-tests were used to test significance of differences between two conditions. When more than one conditions were compared, a one-way ANOVA followed by Dunnett’s multiple comparisons test was performed. A two-way ANOVA followed by Dunnett’s multiple comparisons test were used in cases where more than two groups were compared with repeated measures. In all xenograft assays, we injected 5-8 week old NSG mice, 5 per condition. Both male and female mice were used. When mice died before the end of the experiment due to opportunistic infections the data from those mice was excluded.

## Results

Metastasis is limited by oxidative stress across different cancer types^1,5^. It has been previously shown that metastasizing melanoma cells rely on NADPH regenerating pathways such as one carbon metabolism and pentose phosphate pathway as part of their antioxidant defense system^5,16^. We have used a previously characterized patient-derived xenograft model of melanoma metastasis to quantify levels of NADPH and NADP^+^ in subcutaneous tumors and metastatic nodules in two different patient derived melanoma xenografts (PDX) with different driver mutations, M405 (NRAS Q61H) and M481 (BRAF V600E)^17^. We find that both NADPH and NADP^+^ are elevated in metastatic nodules in different organs compared to the primary tumor in both PDX models (Fig 1 A and B left and middle). The ratio of NADPH/NADP^+^ is an indicator of oxidative stress levels, where a low NADPH/NADP^+^ ratio indicates high levels of oxidative stress in the cells. We have found that the ratio of NADPH/NADP^+^, was decreased in metastatic nodules compared to the subcutaneous tumor in both PDX models, indicating higher levels of oxidative stress (Fig 1 A and B right). Overall, these data suggest that while metastases are under increased levels of oxidative stress, they increase regeneration of NADPH from NADP^+^ as well as *de novo* biosynthesis of NADP^+^ to sustain increased antioxidant capacity. We have also found increased levels of both NAD^+^ and NADH in metastatic nodules compared to subcutaneous tumors across two different PDX models of melanoma metastasis, however the NAD^+^/NADH ratio remained unchanged (Supp Fig 1B and C).

**Figure 1:**
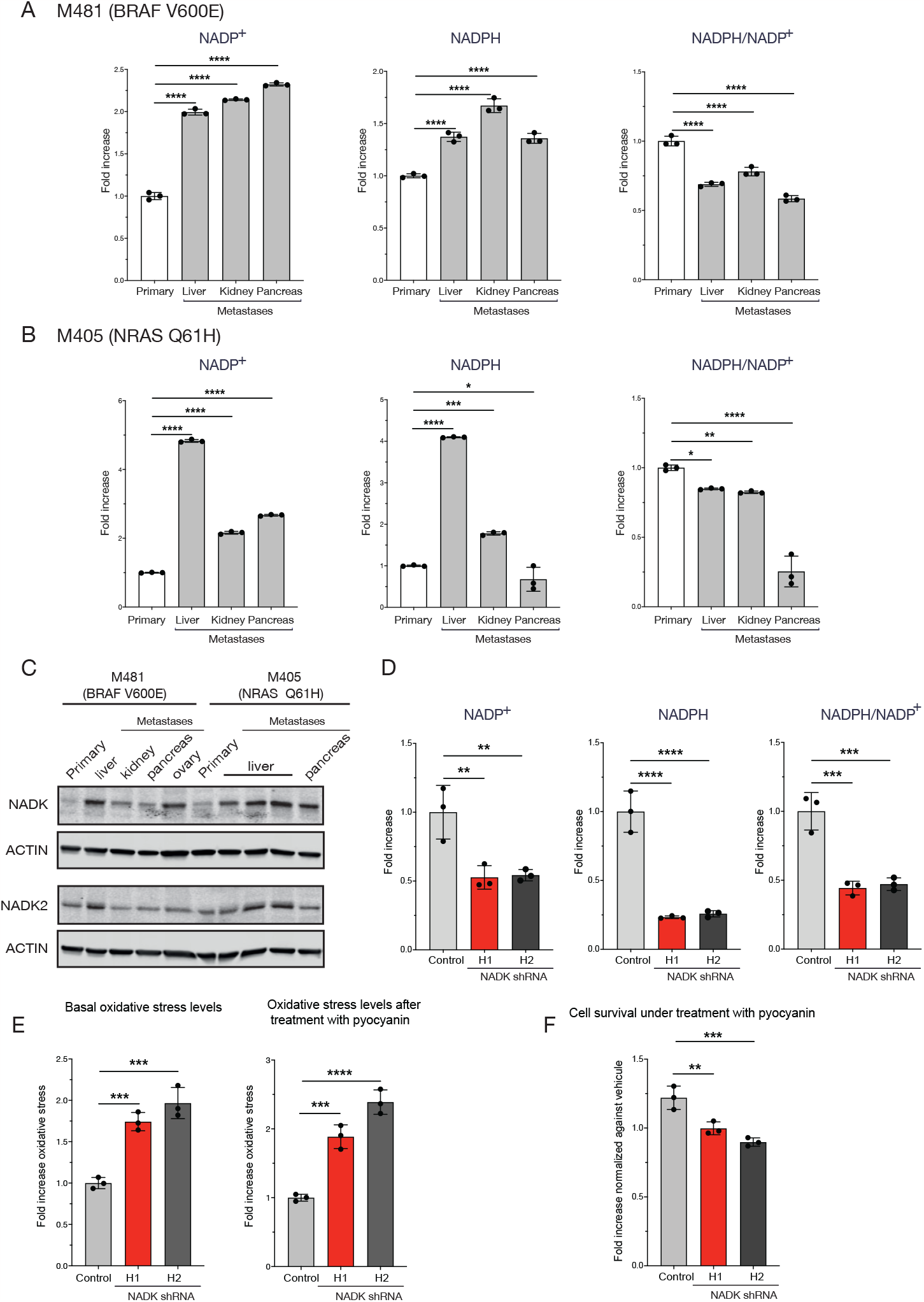
Metastatic nodules exhibit increased synthesis of NADP and NADPH as well as upregulation inase. (A) Levels of NADP^+^ (left), NADPH (middle) and ratio of NADPH/NADP^+^ (right) in subcutaneous melanoma tumors compared to metastatic nodules from different organs in PDX tumor M481. (B) Levels of NADP^+^ (left), NADPH (middle) and ratio of NADPH/NADP^+^ (right) in subcutaneous melanoma tumors compared to metastatic nodules from different organs in PDX tumor M405. (C) Expression levels of NAD^+^ Kinase and NAD^+^ Kinase 2 in subcutaneous melanoma tumors compared to metastatic nodules in using PDX melanoma tumors M481 and M405. (D) Levels of NADP^+^ (left), NADPH (middle) and ratio of NADPH/NADP^+^ (right) in NADK knock down A375 cells. (E) Levels of oxidative stress measured by flow cytometry using Cell ROX Green dye in basal conditions (left) and after treatment with the prooxidant pyocyanin (right) in NADK knock down A375 cells. (F) NADK knock down A375 cells survival after 24hr treated with 10uM pyocyanin.

Since the main mechanism of *de novo* NADP^+^ biosynthesis is through NAD^+^ Kinase (NADK) activity, we tested expression levels of both cytoplasmic NADK and mitochondrial NADK2 in metastatic nodules and subcutaneous tumors in two different PDX models of melanoma metastasis (Fig 1C). Metastatic nodules showed a significant increase in cytoplasmic NADK levels but no significant increase in mitochondrial NADK2 levels, indicating increased NADP^+^ biosynthesis in the cytoplasm during metastasis. We genetically depleted NADK in A375 melanoma cells using shRNAs targeting all three NADK isoforms (Supp Fig 1D Western blot). Levels of both NADP^+^ and NADPH were decreased (Fig 1D left and middle); the ratio of NADPH/NADP^+^ also decreased indicating that these cells are under higher levels of oxidative stress (Fig 1D right). In addition, we found higher levels of Reactive Oxygen Species (ROS) by flow cytometry in NADK-depleted cells, further confirming higher levels of oxidative stress at steady state (Fig 1E left). Loss of NADK lead to higher levels of ROS induced by prooxidant pyocyanin (Fig 1E right) and decreased cell survival, indicating increased sensitivity to oxidative stress (Fig 1F). No significant differences were found in the levels of NAD^+^, NADH and NAD^+^/NADH ratio (Supp Fig 1D graphs).

To functionally test the role of total NADK *in vivo* we knocked down NADK in two PDX melanoma tumors and injected them subcutaneously. There was no difference in the growth of the subcutaneous tumors and a trend towards a decrease in the metastatic burden (Supp Fig 1E and F). We speculate that incomplete depletion of NADK by shRNAs leaves sufficient level of the enzyme to control levels of oxidative stress in order to metastasize.

NADK has three different isoforms^14^; Isoform 1 contains a long autoinhibitory N-terminal domain, while Isoform 3 has a different transcriptional start site and has a unique N-terminal domain (Supp Fig 2A). Due to the difference in the N-terminus domains, Isoform 3 has been previously shown to have a much higher activity than Isoform 1^14^. We tested the mRNA levels of the different isoforms in metastatic nodules compared to the primary tumor in 2 different PDX models of melanoma metastasis. We found that Isoform 3 was greatly upregulated in metastatic nodules compared to Isoform 1 suggesting that the observed increase in NADP^+^ biosynthesis in metastasis is due to increased activity of Isoform 3 (Fig 2A).

**Figure 2:**
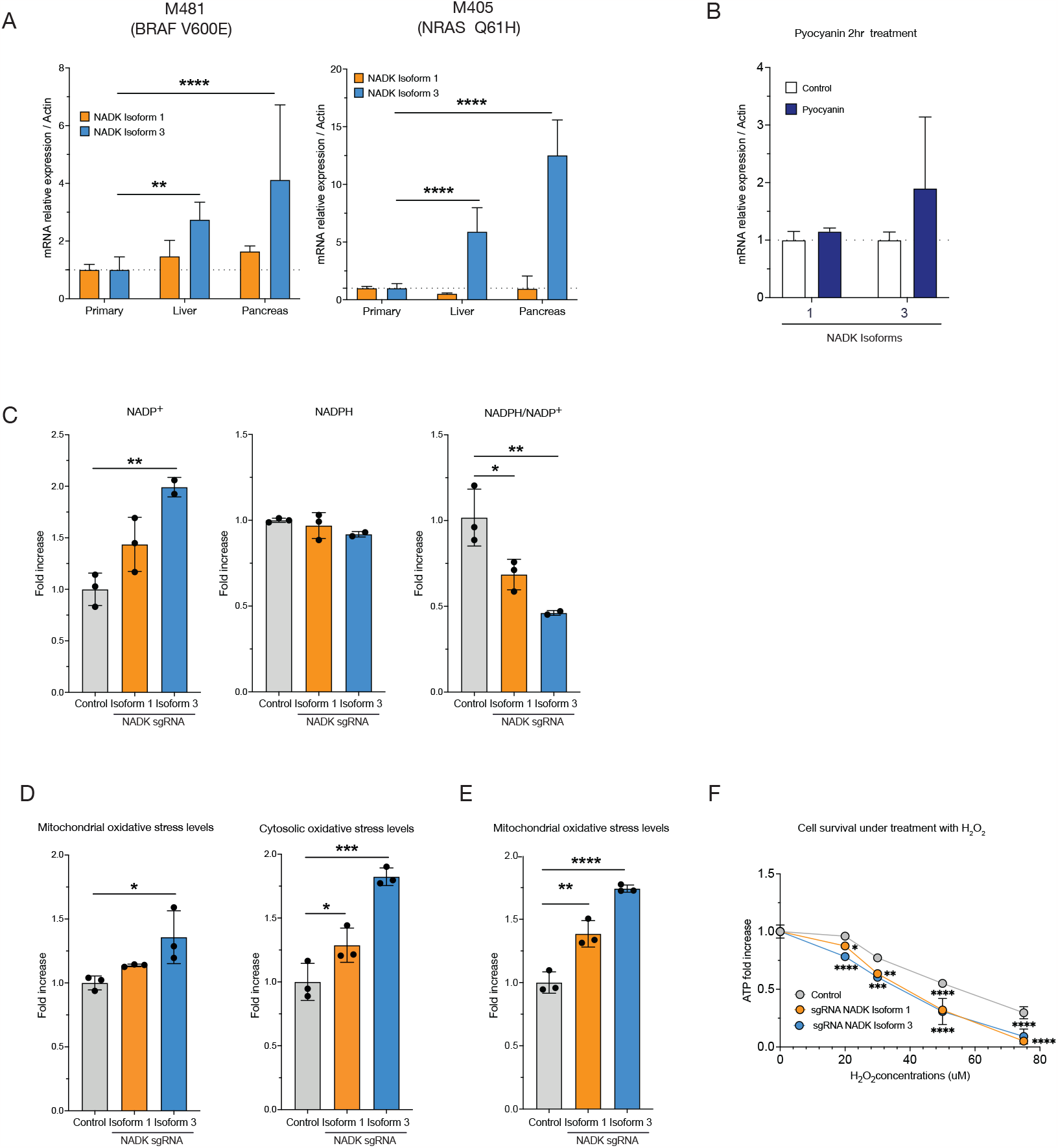
Loss of NADK Isoform 3 increases oxidative stress levels in melanoma cells. (A) mRNA levels of Isoform 1 and 3 in PDX tumor M481 (left) and M4051 (right) in subcutaneous tumor compared to metastatic nodules. (B) NADK Isoform 3 mRNA levels after treatment with 10uM pyocyanin for 2hr. (C) Levels of NADP^+^ (left), NADPH (middle) and ratio of NADPH/NADP^+^ (right) in A375 melanoma cells with Isoform 1 or Isoform 3 deletion. (D) Levels of mitochondrial and cytoplasmic oxidative stress levels by flow cytometry in A375 melanoma cells with either Isoform 1 or Isoform 3 deletions. (E) Mitochondrial oxidative stress levels in A375 melanoma cells by flow cytometry with either Isoform 1 or Isoform 3 deletion treated with 50uM pyocyanin for 2 hrs. (F) Survival of A375 melanoma cells with either Isoform 1 or Isoform 3 deletion under oxidative stress from H_2_O_2_ treatment for 24 hr.

To establish the role of Isoform 3 in oxidative stress regulation, we induced oxidative stress using a prooxidant pyocyanin (Supp Fig 2B) and we saw a transcriptional increase in Isoform 3 but no effect on Isoform 1 (Fig 2B). This suggests that Isoform 3 is specifically upregulated in response to oxidative stress. To establish how the loss of individual NADK isoforms affects melanoma cells we used CRISPR/Cas9 technology to generate isoform-specific knockout A375 melanoma cell lines. Isoform 1 knockout cells were validated by western blotting (Supp Fig 2C left), however due to lack of Isoform 3-specific antibodies, genomic DNA sequencing was used to validate Isoform 3 knockouts (Supp Fig 2C right). Interestingly, we saw an upregulation of Isoform 1 in the Isoform 3 deleted cells suggesting that melanoma cells may compensate for the loss of Isoform 3 with Isoform 1 upregulation. Measurement of NADP^+^ showed an increase in Isoform 3 knockout cells, confirming that there is a compensation with Isoform 1 expression due the lack of Isoform 3 (Fig 2C left). The levels of NADPH showed no change in both knockouts but the NADPH/ NADP^+^ ratio showed a greater decrease in cells lacking Isoform 3 (Fig 2C middle and right), suggesting higher levels of oxidative stress upon loss of Isoform 3. Consistent with decreased NADP^+^ biosynthesis, we found increased levels of both NAD^+^ and NADH in cells lacking either Isoform 1 or Isoform 3 (Supp Fig 2D left and middle). However, we did not find any changes in the NAD/NADH ratio (Supp Fig 2D right). We assessed levels of oxidative stress by flow cytometry using a ROS reactive dye and we found that knockout of Isoform 3 specifically lead to a higher increase in both mitochondrial and cytosolic ROS levels than Isoform 1 knockout (Fig 2D), suggesting that the endogenous compensation by Isoform 1 is not enough to rescue the ROS levels. Oxidative stress induced by prooxidant pyocyanin was also higher in cells lacking Isoform 3 than Isoform 1 as measured by flow cytometry (Fig 2E). We measured the ability of knockout cells to survive under oxidative stress by treating them with the prooxidant H_2_O_2_ and found that both Isoform 1 and 3 knockouts showed decreased cell survival (Fig 2F). Taken together, these data show that loss of Isoform 3 greatly elevates cellular oxidative stress through decreased NADPH/ NADP^+^ ratio and sensitizes melanoma cells to oxidative stress compared to Isoform 1 loss. It also indicates that melanoma cells depend on both isoforms for oxidative stress resistance with Isoform 3 having a greater effect.

Since we observed differential expression of NADK isoforms in metastasis, we wanted to functionally test the role of NADK isoforms in enabling survival of metastasizing melanoma cells during organ colonization *in vivo*. We transplanted luciferase-labelled A375 melanoma cells lacking either NADK Isoform 1 or Isoform 3 subcutaneously into immunocompromised mice. We evaluated metastatic burden in the organs of the mice using bioluminescence signal once the subcutaneous tumor reached the end point of 6 weeks. We found that subcutaneous tumor growth was not affected by either Isoform 1 or Isoform 3 loss (Fig 3 A). However, metastasis was greatly reduced by loss of either Isoform 1 and Isoform 3 suggesting that both isoforms are involved in metastatic colonization (Fig 3B).

**Figure 3:**
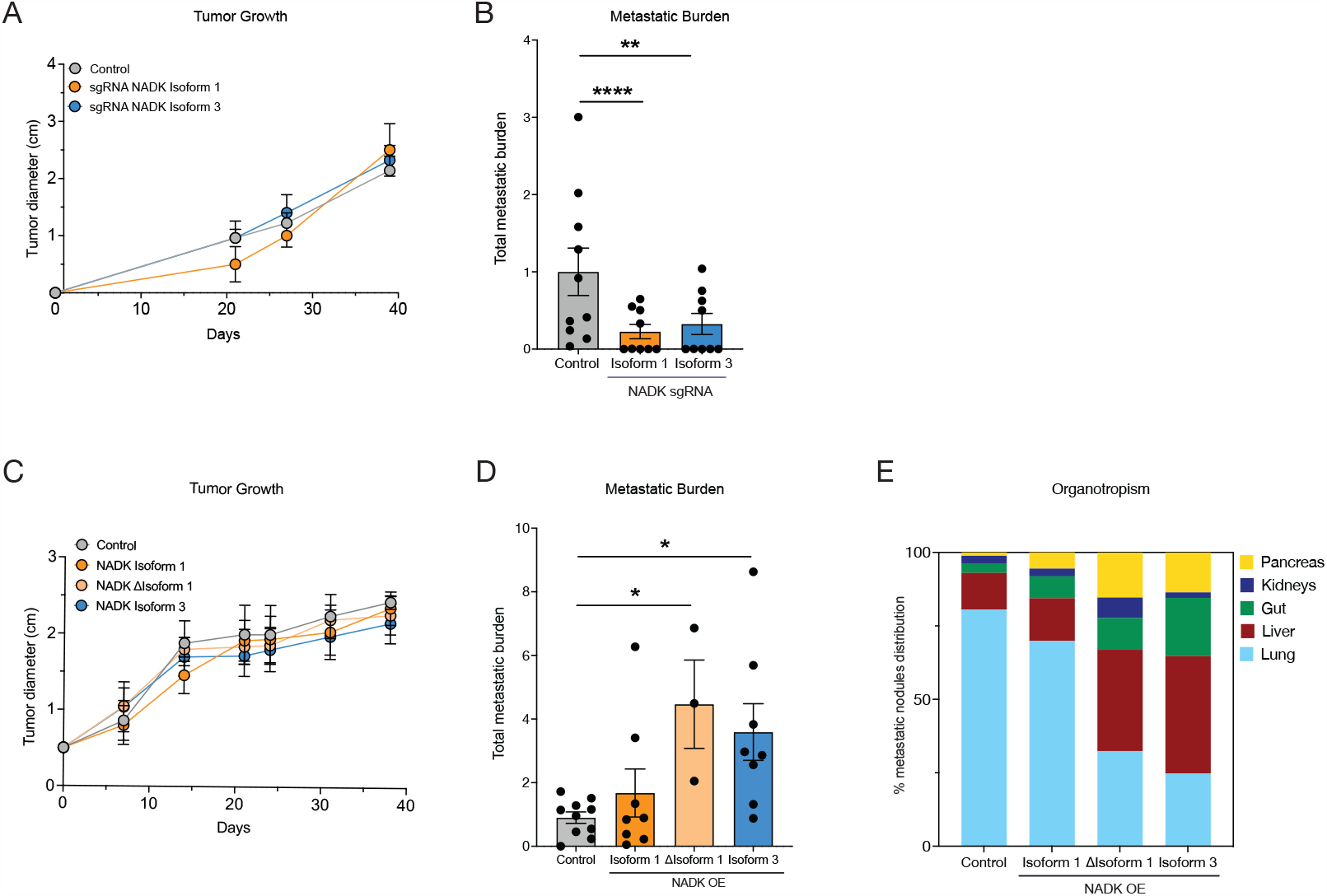
NADK Isoform 3 is neccessary and sufficient for metastasis *in vivo*. (A) Subcutaneous tumor growth of A375 melanoma cells with either Isoform 1 or Isoform 3 deletion in NSG mice. (B) Metastatic burden in the organs of NSG mice xenografted with A375 melanoma cells with either Isoform 1 or Isoform 3 deletion. (C) Subcutaneous tumor growth of A375 melanoma cells overexpressing different NADK isoforms in NSG mice. (D) Metastatic burden in the organs of the metastatic nodules in NSG mice xenografted with A375 melanoma cells overexpressing different NADK isoforms. (E) Organ distribution of the metastatic nodules in NSG mice xenografted with A375 melanoma cells overexpressing different NADK isoforms.

To test the role of the NADK isoforms in our PDX model M405 (NRAS Q61H) we generated an isoform specific knockdown, that we verified by flow cytometry using an isoform specific reporter in A375 melanoma cells (Supp Fig 3A and B). We injected PDX melanoma cells with either Isoform 1 or Isoform 3 knockdown subcutaneously and measured both primary tumor growth and metastatic burden. At the end point of the experiment we found no difference in the growth of the primary tumor between control and Isoform 1 and a slight decrease in Isoform 3 (Supp Fig 3C left), but there was a significant decrease in metastatic burden upon loss of either isoform (Supp Fig 3C right).

In order to determine whether NADK isoforms can differentially drive metastasis *in vivo*, we overexpressed NADK isoforms 1 and 3 in Luciferase labelled A375 melanoma cells. In addition, we overexpressed a truncated version of Isoform 1,(∆Isoform 1) (Supp Fig 2A and Suppl Fig 3D). Trunctated Isoform 1 lacks the regulatory N-terminus domain and it has been shown to have a similar activity level as Isoform 3 *in vitro*^14^. We tested the levels of NADP^+^, NADPH and the ratio NADPH/NADP^+^ in these cells, and as expected Isoform 3 and ∆Isoform 1 had the highest levels of NADP, NADPH and NADP/NADPH ratio (Supp Fig 3E)

We transplanted A375 cells subcutaneously and we found that none of the isoforms had a significant impact on the growth of the subcutaneous tumor (Fig 3C). Overexpression of wild type Isoform 1 had no effect on the amount of metastatic burden in the organs of the mice, however, overexpression of either truncated Isoform 1 or Isoform 3 significantly increased the metastatic burden in the organs of the mice (Fig 3D). This suggests that overall increased *de novo* biosynthesis of NADP^+^ drives metastasis. When we analyzed the metastatic organotropism of melanoma cells overexpressing different isoforms, we found that both truncated Isoform 1 and Isoform 3 showed significantly less metastasis to the lung and more metastasis to other organs such as the liver, pancreas and the gut, compared to Isoform 1-overexpressing cells and control cells (Fig 3E). This suggests that increased *de novo* NADP^+^ biosynthesis by truncated Isoform 1 or Isoform 3 drives broad metastatic colonization to different organs. This data also suggests that cells with low NADP^+^ biosynthesis may get lodged in the lung and are unable to metastasize more widely to other sites.

Since NRF2 is the major transcriptional regulator of the antioxidant response we analyzed existing ChIP data from the ENCODE database and found that NRF2 was specifically found to bind the putative promoter of Isoform 3 but not Isoform 1 in Hela-S3 cells (Fig 4A) and A549 cells (Data not shown). The NFR2 binding sequence also known as antioxidant response element (ARE) was found in the Isoform 3 promoter (Fig 4B). To test this hypothesis functionally, we used NRF2 activator Ki696 to test its effect on Isoform 3 induction in A375 melanoma cell line. We found that NRF2 activation leads to a rapid increase in Isoform 3 level, but not in Isoform 1 (Fig 4C and Supp 4A). We generated NRF2 knockdown in A375 using shRNA (Supp Fig 4B and C). The levels of Isoform 3 were significantly reduced after treatment of the knockdown cells with the NRF2 inducer, Ki696. We also found an increase of Isoform 1, suggesting again that there is a compensation between the two isoforms (Fig 4D). This data suggests that Isoform 3 is upregulated in response to oxidative stress, specifically through the action of NRF2, and that in the absence of Isoform 3 the cell compensates by inducing the expression of Isoform 1, confirming the importance of Isoform 3 in the antioxidant response.

**Figure 4:**
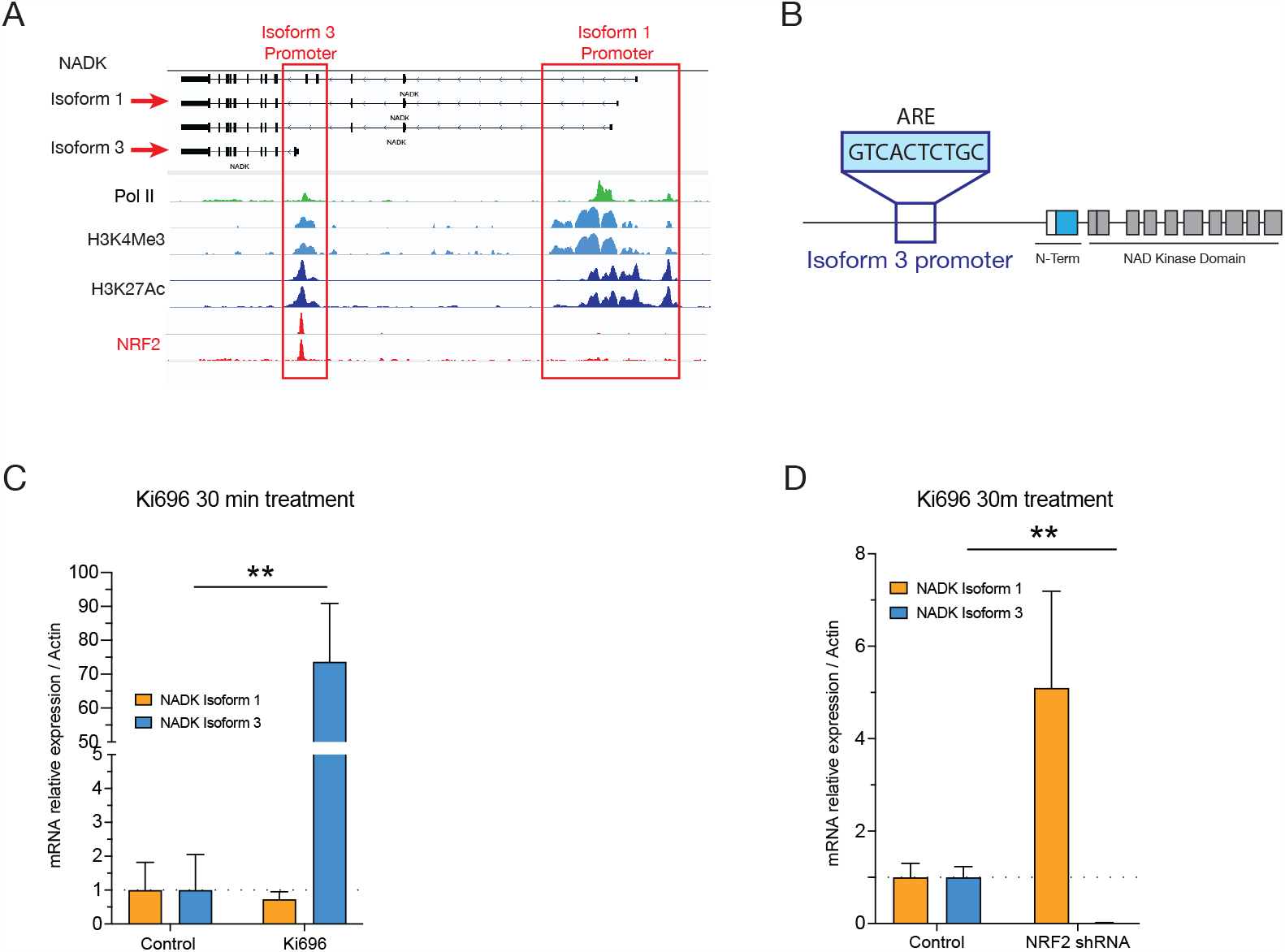
NRF2 specifically regulates NADK Isoform 3. (A) ChIP seq data looking at Pol2 and activating histone marks. (B) Schematic of the sequence of the ARE in the Isoform 3 promoter. (C) NADK Isoform 1 and 3 mRNA expression in A375 melanoma cell line treated with Ki676 (5uM) for 30 minutes. (D) NADK Isoform 1 and 3 mRNA expression in A375 melanoma cell line NRF2 knock down treated with Ki676 (5uM) for 30 minutes.

## Discussion

Metastasizing cancer cells have increased plasticity and are able to rewire their metabolism to increase survival during metastatic colonization. Increased antioxidant capacity enables metastasis of cancer cells by increasing regeneration of GSH from GSSG through use of NADPH and its consequent conversion to NADP^+^. While there are numerous metabolic pathways that can recycle NADPH from NADP^+^ to drive the cellular antioxidant cycle, such as the pentose phosphate pathway, malic enzyme and one carbon metabolism, there are limited ways that the NADP^+^ pool can be replenished *de novo*. NAD kinase (NADK) is the main route of generating NADP^+^ from NAD^+^. Since NADPH is used for reduction reactions, cells maintain a high NADPH/ NADP^+^ ratio and therefore convert most of NADP^+^ into NADPH. Increased activity of NADK would contribute to increased antioxidant capacity of cancer cells, through *de novo* synthesis of NADP^+^ and therefore generation of NADPH for GSH regeneration.

It has been previously shown that NADK has a role in cancer and metastasis. In breast cancer NADK levels are increased due to a histone H3.3 variant-mediated epigenetic regulation of the NADK promoter helping the cancer cells to metastasize^18^. An increased occurrence of mutant NADK (NADK^I90F^) with increased activity has been found to increase cancer cell growth in pancreatic adenocarcinoma^19^. Consistent with these findings, we have found higher levels of NADK in metastatic nodules compared with the primary tumor in a patient-derived xenograft model of melanoma. Increased expression of NADK would increase the supply of NADP^+^ that is reduced to NADPH allowing cells to survive under high levels of oxidative stress. We have shown loss of NADK makes melanoma cells more sensitive to prooxidant treatment and less metastatic.

NADK kinase has been shown to exist in several isoforms. Isoforms 1 has a long-term autoinhibitory N-terminus domain, whose inhibition is alleviated upon post-translational phosphorylation. One of the major regulators of NADK activity has been shown to be AKT signaling as well as PKC signaling^14,15^. Both kinases have been shown to phosphorylate specific sites in the autoinhibitory N-terminus, which means that increased signaling through the AKT and PKC pathways leads to increased NADK activity and increased NADP^+^ production. In contrast, Isoform 3 of NADK has a unique short N-terminal domain, generated by an alternative transcriptional start site, lacking autoinhibitory properties and remaining outside of upstream kinases regulation. This also means that Isoform 3 has a much higher activity than the other Isoforms at baseline. AKT signaling through AKT1 has been shown to decrease during epithelial-mesenchymal transition^20,21^. In fact, genetic depletion of AKT1 has been shown to promote an invasive phenotype in breast cancer^20^. This suggests that there may not be sufficient AKT signaling to alleviate the autoinhibition of NADK Isoform 1 during metastasis, however metastasizing cells are still highly dependent on the NADP(H) pool for antioxidant defenses. Our findings suggest that Isoform 3 due to its high activity, which is independent of AKT signaling, is a potentially novel mechanism to increase the NADP(H) pool and increase oxidative stress resistance.

Our results *in vitro* reveal that Isoform 3 expression increases in response to oxidative stress and it is highly upregulated in metastatic nodules. Lack of both isoforms in melanoma cells increases levels of oxidative stress having a higher impact in Isoform 3 knock out cells. Our data suggest that both Isoform 1 and 3 are required for metastatic colonization, while loss of either isoform does not have an impact on the primary tumor growth. This may indicate that the demand for NADPH is much higher in metastasizing melanoma cells compared to proliferating cells in the primary tumor. This also suggests that even a minor decrease in NADP^+^ levels has a strong effect on the ability of melanoma cells to colonize distant sites.

Interestingly, both truncated Isoform 1 and Isoform 3 overexpression led to an increase in the metastatic burden in the organs of the mice, while Isoform 1 overexpression had no effect. Since it has been previously shown that truncated Isoform 1 and Isoform 3 have similar activities *in vitro*, and they both increase metastatic colonization, this data suggests that increased *de novo* NADP^+^ generation is a major metastatic driver. Intriguingly, when we analyzed metastatic organotropism we saw increased metastasis to the liver, gut and pancreas and decreased metastasis to the lung with cells overexpressing truncated Isoform 1 and Isoform 3. On the other hand, control cells and full-length Isoform 1 overexpressing cells showed increased metastasis to the lung and limited metastasis to other organs. This suggests that during metastasis, first many cells get lodged in the lung and tend to metastasize there, however increased NADP^+^ biosynthesis enables cells to metastasize more widely. It is possible that after dissemination, cells with increased NADP^+^ biosynthesis are able to survive better across many different organs, and therefore cause more widespread metastasis. More broadly, this indicates that there are different mechanisms of oxidative stress resistance that are required for melanoma cells at different steps of the metastatic cascade and in different organotropic environments.

Metastasizing cancer cells experience many different stresses during the metastatic cascade. It is likely that these stress cues induce specific stress resistance mechanisms. Our work specifically identified NRF2, a master regulator of the antioxidant response as the transcription factor responsible for driving Isoform 3 expression. Our data suggests that Isoform 3 is rapidly upregulated by NRF2. This suggests that Isoform 3 may play a role in the initial response to acute oxidative stress due to its elevated activity, while Isoform 1 may play a role in the maintenance of oxidative stress resistance, suggesting there are different mechanisms between acute and chronic oxidative stress resistance during metastasis. Upregulation of Isoform 3 maybe part of a broader network of oxidative stress adaptations that occur during metastatic colonization.

In conclusion, our work has discovered an antioxidant mechanism through differential transcriptional induction of various NADK isoforms. We find Isoform 3 is specifically upregulated in metastatic nodules compared to the primary tumor and provides antioxidant protection to melanoma cells increasing extravasation and organ colonization by metastasizing melanoma cells, driven by coordinated action of NRF2. Taken together, our data highlights NADK Isoform 3 as a potential novel mechanism that drives metastatic colonization that could be exploited for therapeutic inhibition of metastatic disease.

## Supplementary Figures

**Supplementary Figure 1.**
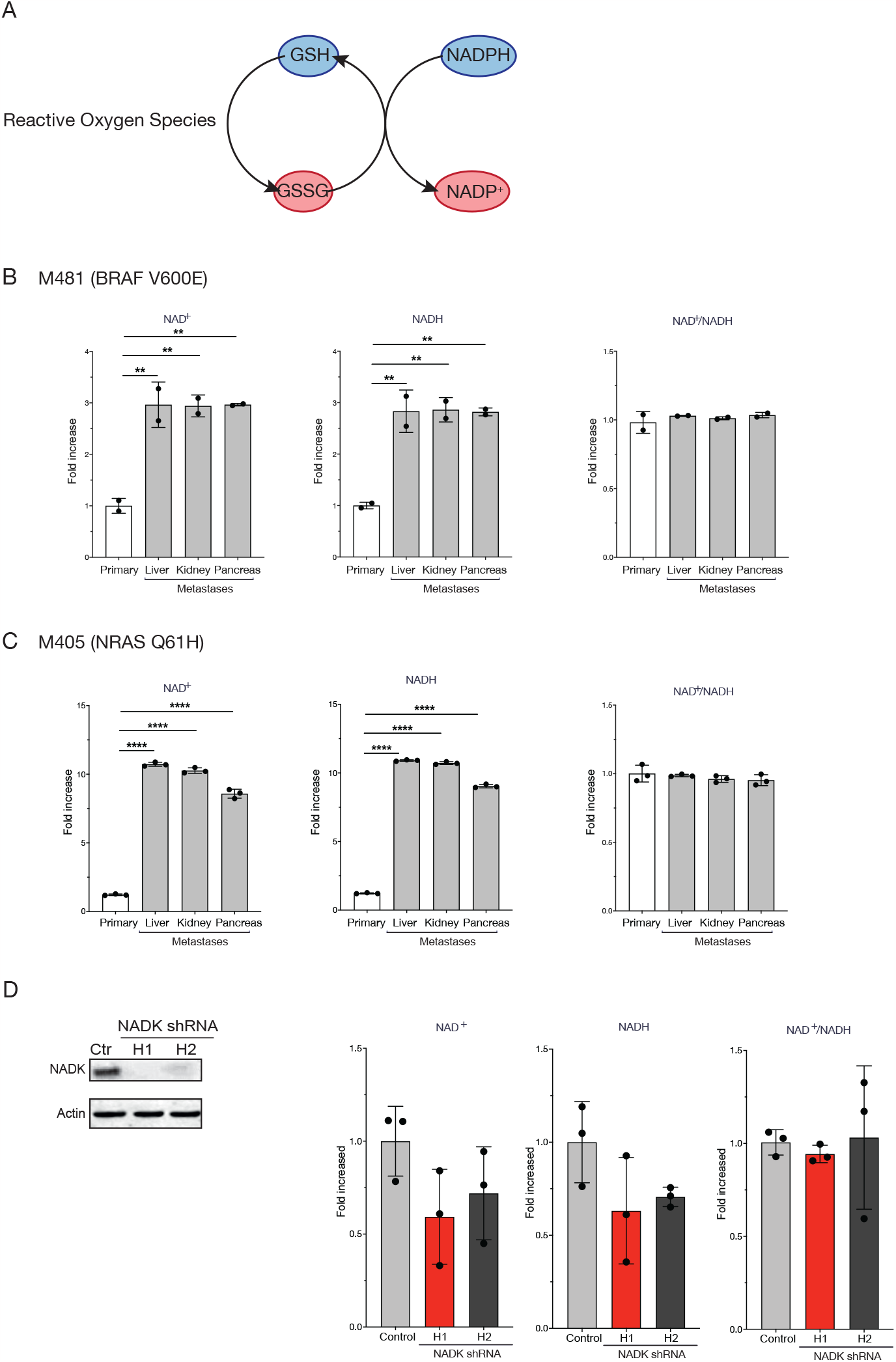
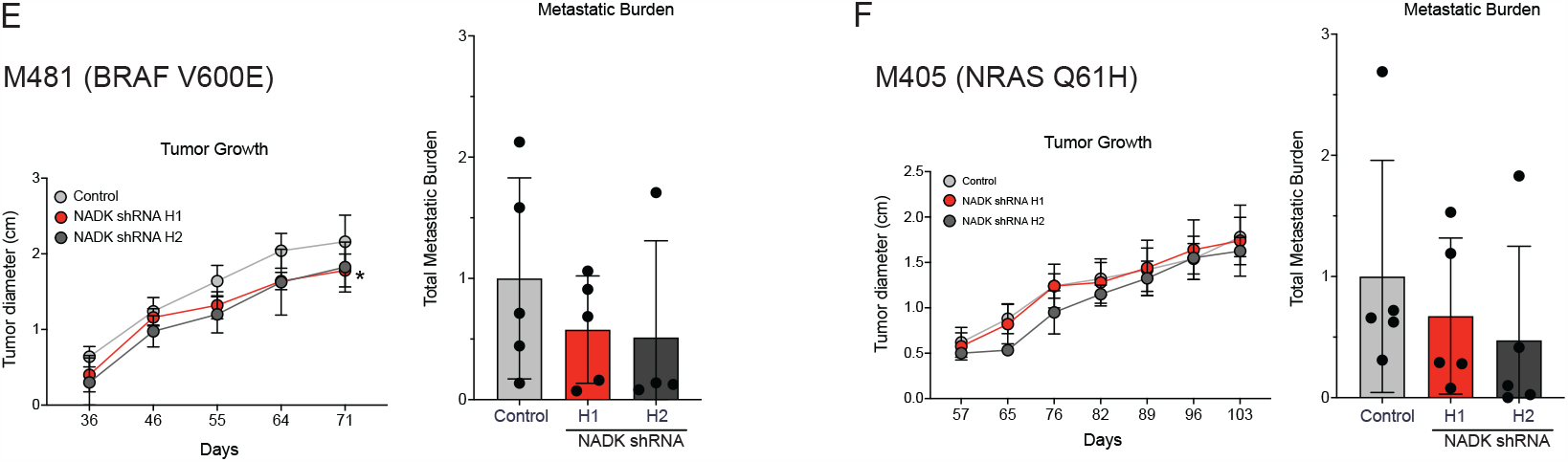
(A) Levels of NAD^+^ (left), NADH (middle) and ratio of NAD^+^/NADH (right) in subcutaneous melanoma tumors compared to metastatic nodules from different organs in PDX tumor M481. (B) Levels of NAD^+^ (left), NADH (middle) and ratio of NAD^+^/NADH (right) in subcutaneous melanoma tumors compared to metastatic nodules from different organs in PDX tumor M405. (C) Expression levels of NADK in NADK knock down A375 cells. (D) Levels of NAD^+^ (left), NADH (middle) and ratio of NAD^+^/NADH in NADK knock down A375 cells. (E) Subcutaneous tumor growth of PDX tumor M481with NADK expression knock down in NSG mice (left). Metastatic burden in the organs of the metastatic nodules in these NSG mice (right). (F) Subcutaneous tumor growth of PDX tumor M405 with NADK expression knock down in NSG mice (left). Metastatic burden in the organs of the metastatic nodules in these NSG mice (right).

**Supplementary Figure 2.**
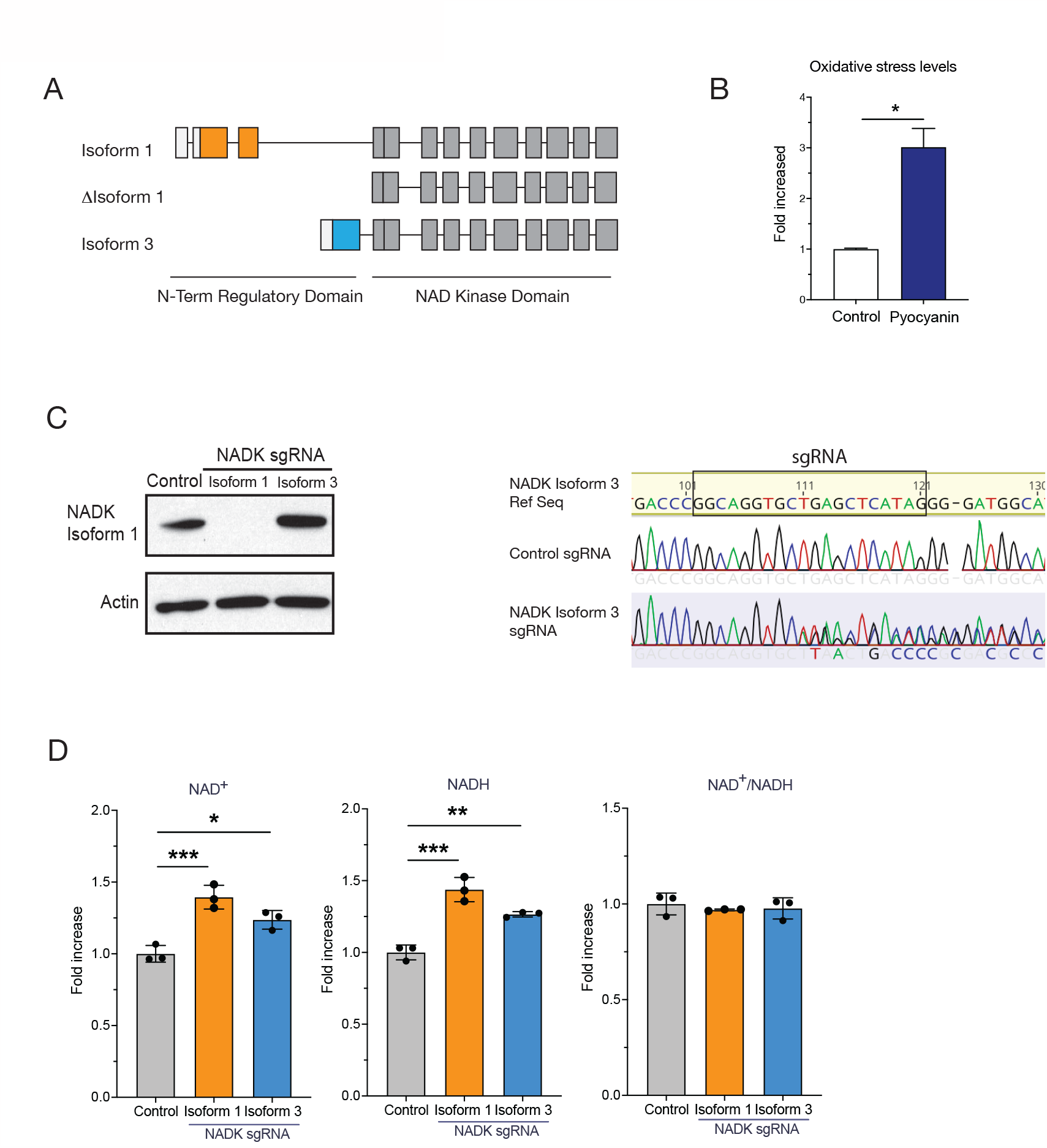
(A) Schematic of NAD^+^ Kinase Isoforms 1, truncated Isoform 1 and Isoform 3. (B) Mitochondrial oxidative stress levels in A375 melanoma cells by flow cytometry treated with pyocyanin (10uM for 2 hr). (C) Western blot for NADK Isoform 1 levels (left) and sequencing results (right) in A375 melanoma cells with Isoform 1 or Isoform 3 deletions. (D) Levels of NAD^+^ (left), NADH (middle) and ratio of NAD^+^/NADH in A375 melanoma cells with Isoform 1 or Isoform 3 deletions.

**Supplementary Figure 3.**
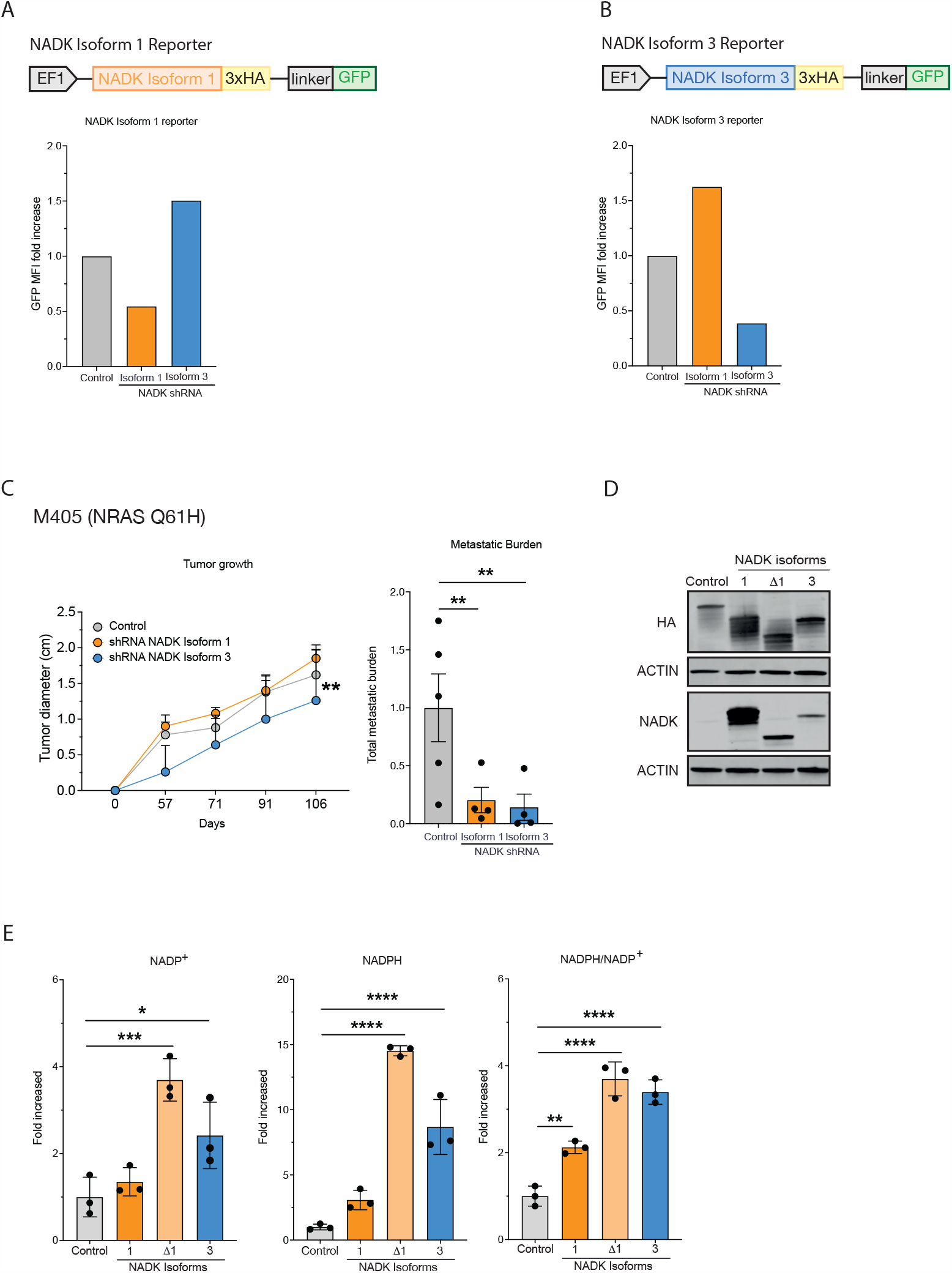
(A) Schematic of NADK Isoform 1 reporter (top). Levels of NADK Isoform 1 by flow cytometry in NADK Isoform 1 and Isoform 3 knock down in A375 cells. (B) Schematic of NADK Isoform 3 reporter (top). Levels of NADK Isoform 1 by flow cytometry in NADK Isoform 1 and Isoform 3 knock down in A375 cells. (C) Left. Subcutaneous tumor growth of A375 NADK Isoform 1 and Isoform 3 knock down in NSG mice. Right. Metastatic burden in the organs of the metastatic nodules of A375 NADK Isoform 1 and Isoform 3 knock down in NSG mice. (D) Western blot with NADK Isoform overexpression cell lines. (E) Levels of NADP^+^ (left), NADPH (middle) and ratio of NADPH/ NADP^+^ in A375 melanoma cells with Isoform 1, truncated Isoform 1 and Isoform 3 deletions.

**Supplementary Figure 4.**
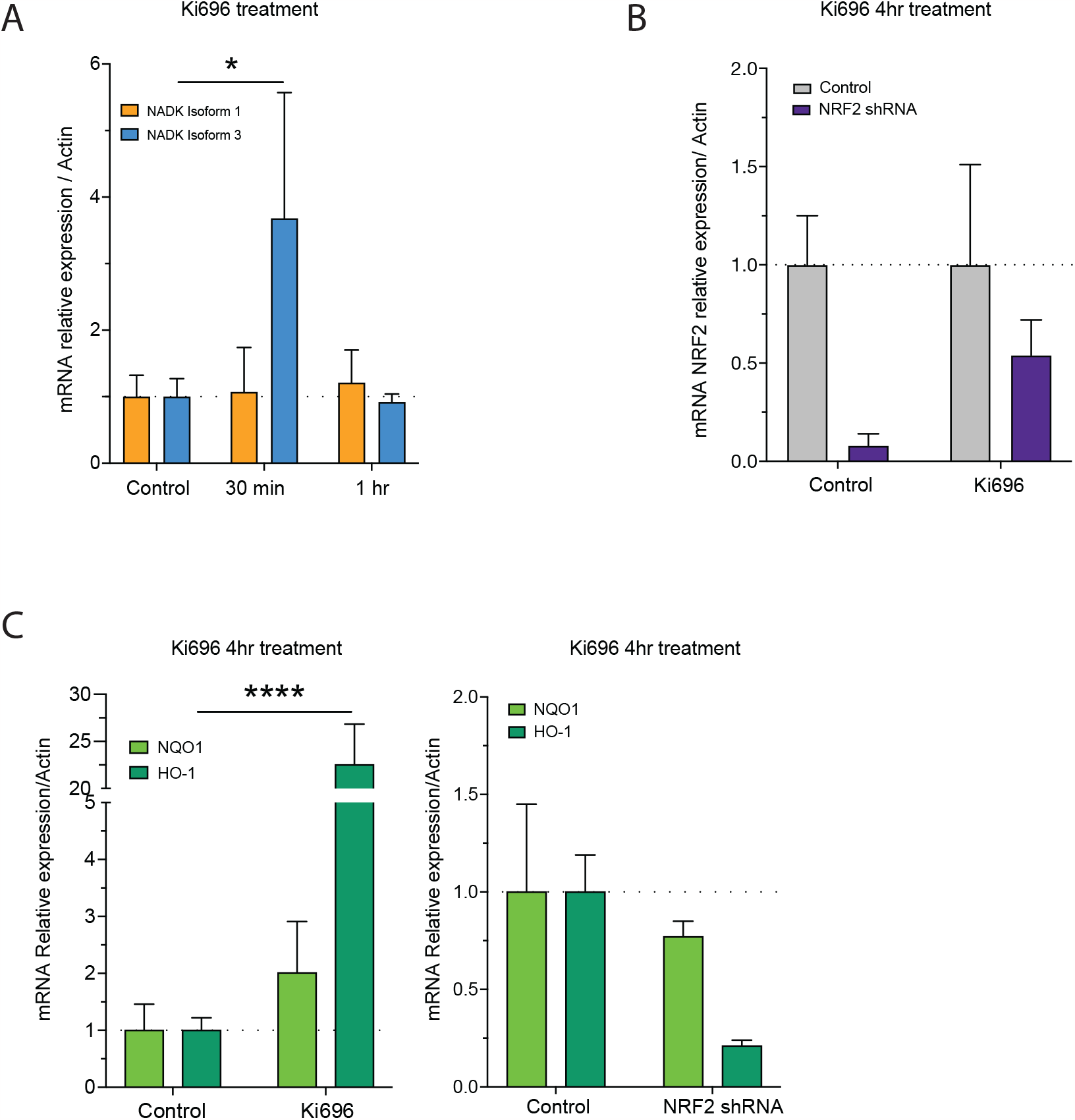
(A) NADK Isoform 1 and 3 mRNA expression in A375 melanoma cell line treated with Ki676 (5uM) for 30 minutes and 1hr. (B) mRNA levels of NRF2 in A375 melanoma cell line NRF2 knock down treated with Ki676 (5uM) for 4hr. (C) mRNA levels of NRF2 targets HO-1 and NQO-1 in A375 melanoma cell line (left) and in A375 melanoma cell line NRF2 knock down (right) after 4hr treatment with Ki696 5uM.

