## Supplementary Figures for "Transcriptional Isoforms of NAD^+^ Kinase regulate oxidative stress resistance and melanoma metastasis"

### Supplementary Figure 1

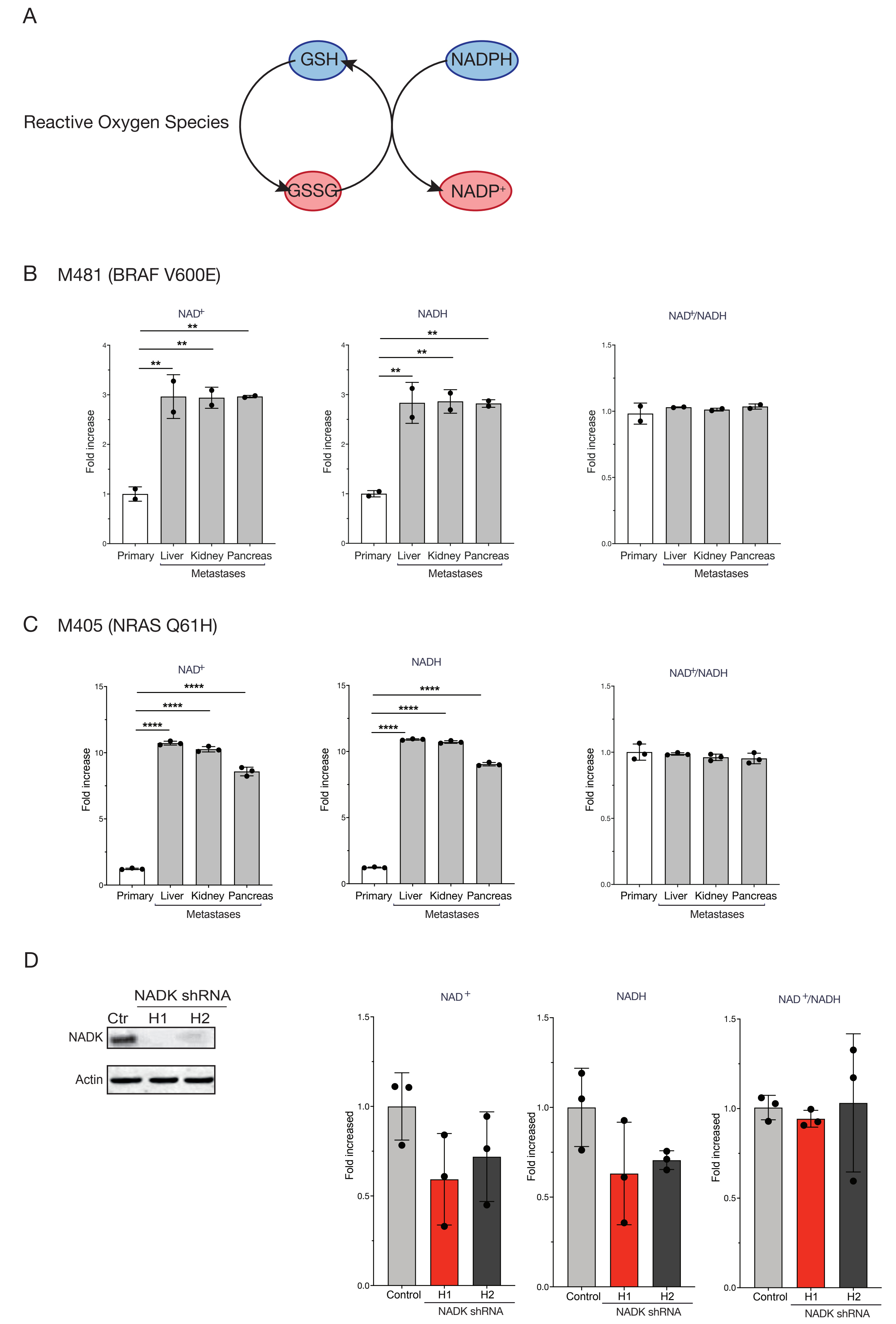

### Supplementary Figure 1

E

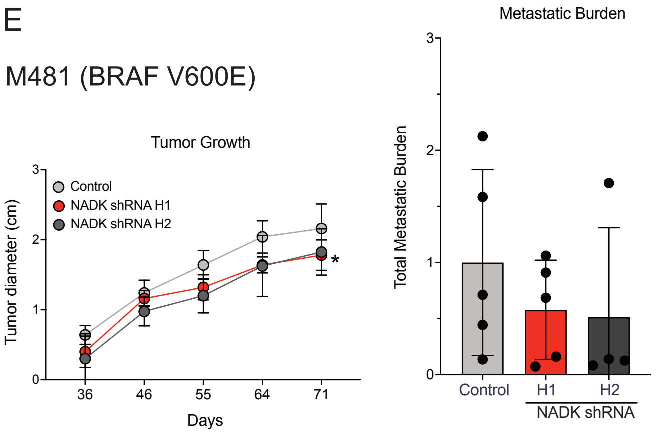


F

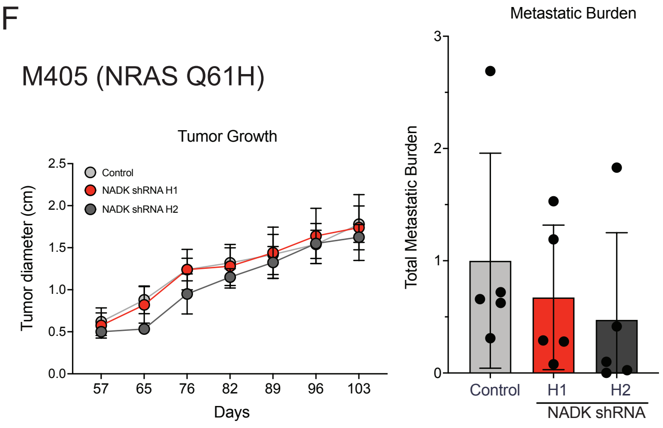

Supplememtary Figure 2

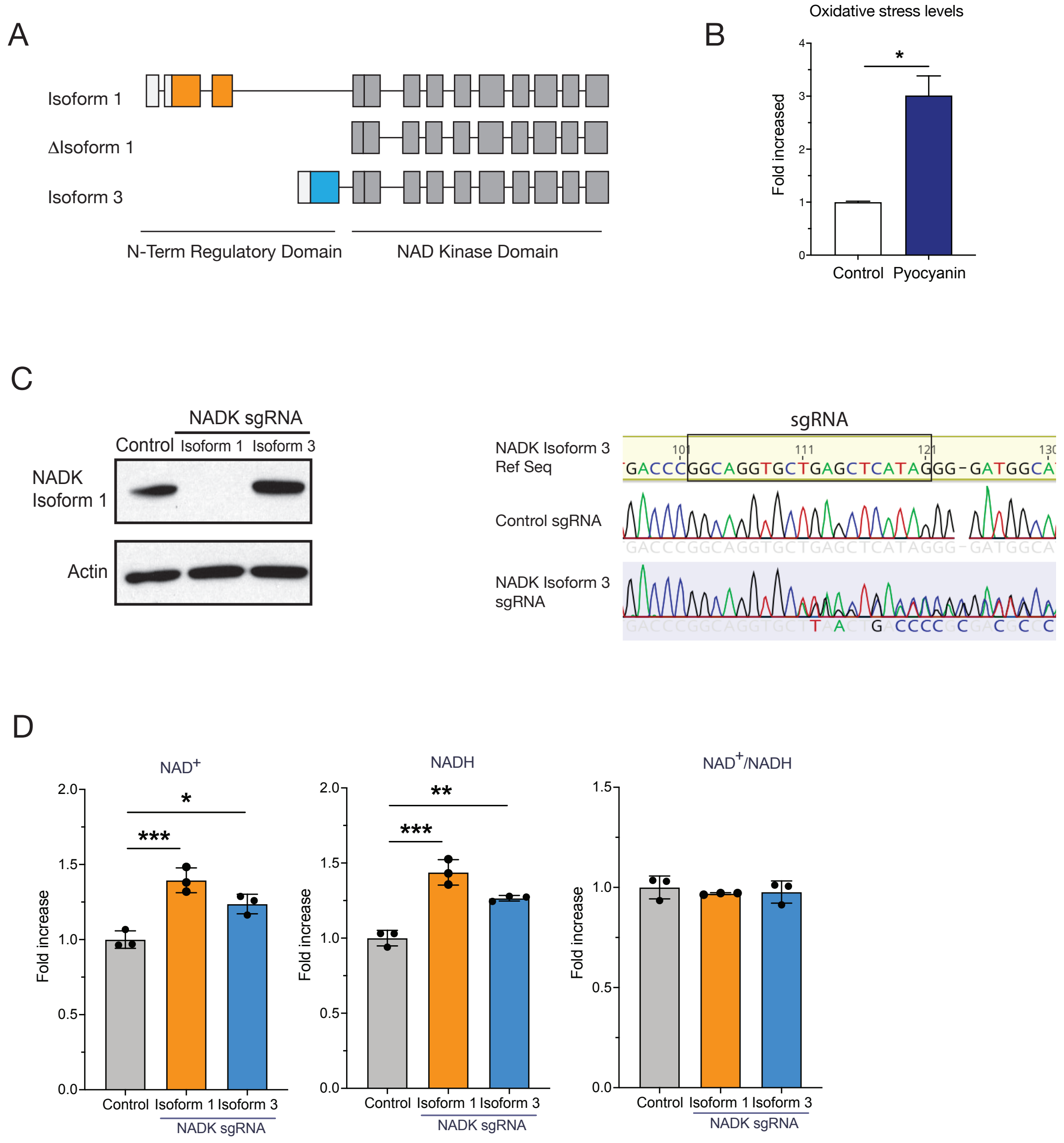

Supplementary Figure 3

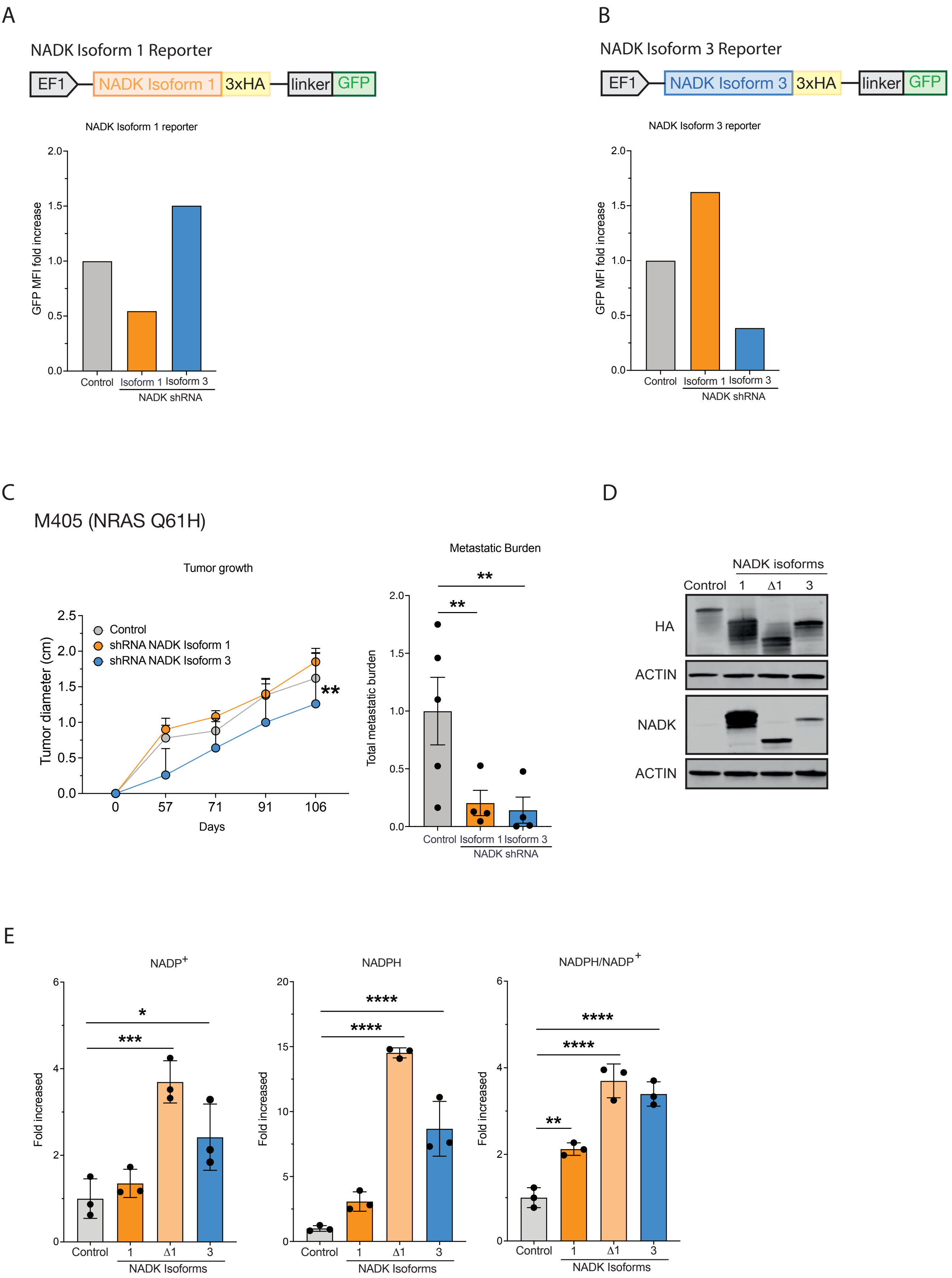

### Supplementary Figure 4

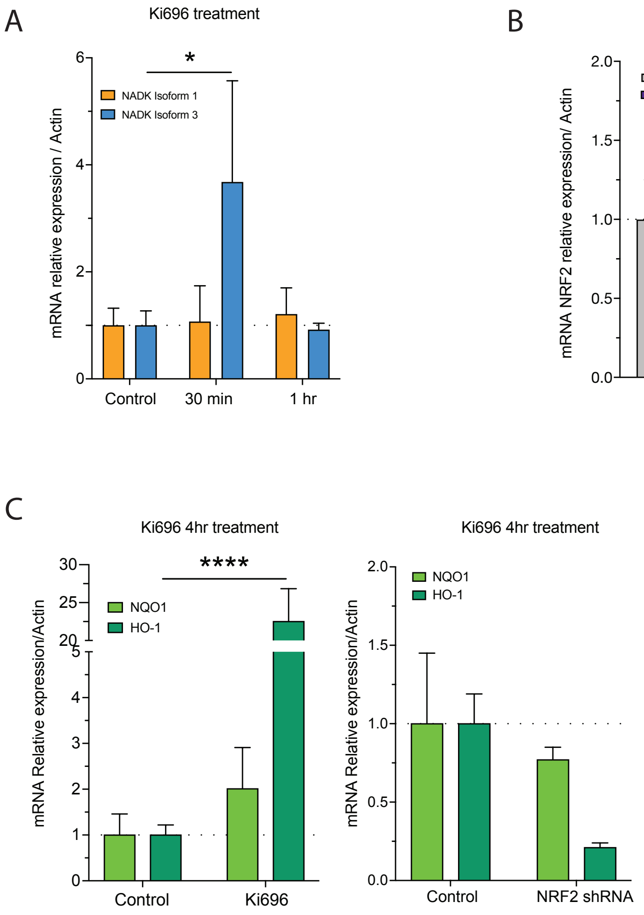
